# Class of antiretroviral drugs and anemia risk in the current treatment era

**DOI:** 10.1101/674549

**Authors:** B.N. Harding, B.M. Whitney, R.M. Nance, H.M. Crane, G. Burkholder, R.D. Moore, W.C. Mathews, J.J. Eron, P.W. Hunt, P. Volberding, B. Rodriguez, K.H. Mayer, M.S. Saag, M.M. Kitahata, S.R. Heckbert, J.A.C. Delaney

**Affiliations:** University of Washington; University of Alabama Birmingham; Johns Hopkins University; University of California San Diego; University of North Carolina, Chapel Hill; University of California San Francisco; Case Western Reserve University; Fenway Health Institute

**Keywords:** antiretroviral agents, cohort, anemia, integrase inhibitors, protease inhibitors, non-nucleoside reverse transcriptase inhibitors

## Abstract

**OBJECTIVES:** Anemia is common among people living with HIV (PLWH) and has been associated with certain, often older, antiretroviral medications. Information on current antiretroviral therapy (ART) and anemia is limited. The objectives were to compare associations between anemia incidence or hemoglobin change with core ART classes in the current ART era.

**DESIGN:** Retrospective cohort study.

**SETTING:** U.S.-based prospective clinical cohort of PLWH aged 18 and above receiving care at 8 sites between 1/2010-3/2018.

**PARTICIPANTS:** 16,505 PLWH were included in this study.

**MAIN OUTCOME MEASURES:** Anemia risk and hemoglobin change were measured for person-time on a protease inhibitor (PI) or an integrase strand transfer inhibitor (INSTI), relative to a non-nucleoside reverse transcriptase inhibitor (NNRTI) reference. We also examined PLWH on multiple core classes. Cox proportional hazards regression analyses were conducted to measure associations between time-updated ART classes and incident anemia or severe anemia. Linear mixed effects models were used to examine relationships between ART classes and hemoglobin change.

**RESULTS:** During a median of 4.9 years of follow-up, 1,040 developed anemia and 488 developed severe anemia during. Compared to NNRTI use, INSTI-based regimens were associated with an increased risk of anemia (adjusted hazard ratio [aHR] 1.17, 95% confidence interval [CI] 0.94-1.47) and severe anemia (aHR1.55 95%CI 1.11-2.17), and a decrease in hemoglobin level. Time on multiple core classes was also associated with increased anemia risk (aHR 1.30, 95%CI 1.06-1.60) and severe anemia risk (aHR 1.35, 95%CI 0.99-1.85), while no associations were found for PI use.

**CONCLUSION:** These findings suggest INSTI use may increase the risk of anemia. If confirmed, screening for anemia development in users of INSTIs may be beneficial. Further research into underlying mechanisms is warranted.

**Strengths and limitations of this study:**

- This study utilized a large and geographically diverse population of PLWH in care across the U.S.
- This study leveraged comprehensive clinical data, including information on diagnoses, medication use, laboratory test results, demographic information, and medical history.
- This study investigated associations between specific types of ART core regimens and anemia risk.
- This observational study is subject to residual confounding.
- This study focused on anemia assessed from hemoglobin lab values taken at regular medical care visits without excluding participants with conditions strongly associated with hemoglobin level through non-traditional HIV mechanisms.

## Introduction

Anemia (hemoglobin [Hb]<10 g/dL) and severe anemia (Hb<7.5 g/dL) [1] are common among people living with HIV (PLWH) [2]. The prevalence of anemia is elevated in PLWH compared to the general population. One study reported that among non-pregnant American women living with HIV, the prevalence of anemia was 28.1% compared to 15.1% among women without HIV [3]. Estimates vary by age, sex, HIV disease stage, use of antiretroviral therapy (ART) and injection drug use status [2, 4]. Among PLWH, associations have been found between anemia and mortality [5–10] health-related quality-of-life [2], morbidity, dementia [11], and treatment failure [12]. In addition, anemia is an independent prognostic indicator associated with HIV disease progression [2, 13, 14], including development of AIDS [8].

Research shows that ART impacts anemia risk among PLWH. In the early treatment era, use of zidovudine (AZT) was a cause of bone marrow suppression leading to anemia [15]. However, in recent years, AZT use has decreased substantially as other, better tolerated ART medications have become available. Despite the impact of specific agents such as AZT, ART use in general is associated with reduced anemia incidence [16, 17] likely due to inhibition of HIV disease progression. Since worsening HIV disease parameters are associated with anemia, better disease control with ART reduces the risk of anemia [18]. Current ART regimens typically include a pair of nucleoside reverse transcriptase inhibitors (NRTIs) as a backbone plus a core agent. Common core classes include non-nucleoside reverse-transcriptase inhibitors (NNRTIs), integrase strand transfer inhibitor (INSTIs), and protease inhibitors (PIs). While ART use overall reduces anemia, little is known about whether anemia risk differs between commonly used ART classes in the current treatment era, particularly the newer INSTI class. Some studies found a possible increased rate of anemia among PI users [19], and in a randomized controlled trial, some participants discontinued INSTI due to anemia adverse events [20]. However, many studies included few participants or were mostly from an earlier ART era when older ART medications were predominantly used. The objective of this study was to compare rates of anemia and severe anemia development as well as changes in Hb overtime based on classes of ART used in the current treatment era.

## Methods

### Overview and setting

The present study included PLWH in care in the Centers for AIDS Research (CFAR) Network of Integrated Clinical Systems (CNICS) cohort during the period of January 1, 2010 to March 31, 2018. The CNICS cohort has been described in detail elsewhere [21]. Briefly, CNICS is a dynamic prospective clinical cohort of >32,000 adult PLWH receiving care at eight participating sites across the U.S. Comprehensive clinical data, including diagnoses, ART and other medications, laboratory test results, demographic information, and historical information, including ART use before enrollment, is collected through electronic medical records and other institutional data systems at each site and harmonized in the CNICS data repository. Medication data including ART use are entered into the electronic medical records by clinicians or prescription fill/refill data are uploaded directly from pharmacy systems and verified through medical record review. Participants entered the current study on January 1, 2010 and the earliest date that they met the following enrollment criteria (cohort entry date): a) enrollment in CNICS for ≥6 months to allow time for covariate ascertainment and b) use of an ART regimen containing a PI, NNRTI, or INSTI. In addition, all participants were required to have at least 2 available hemoglobin lab values during study follow-up. Figure 1 shows inclusion criteria and exclusions made. Informed consent was obtained from all participants and institutional review boards at each site approved CNICS protocols.

**Figure 1:**
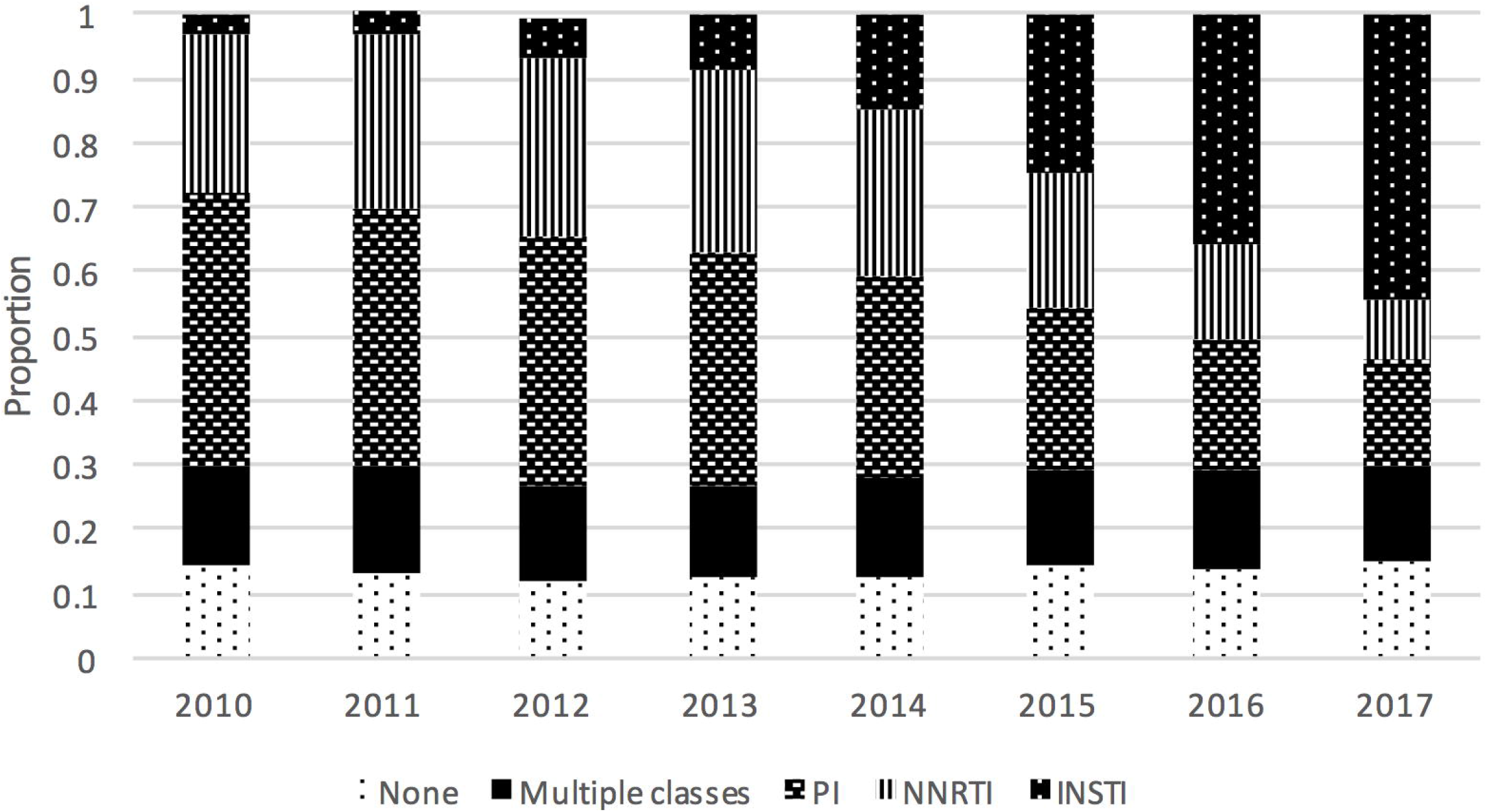
Flow chart for inclusion/exclusion criteria for 22,027 PLWH in care at CNICS after 1/2010. Exclusions were made for those not exposed to any of the ART core classes, those with fewer than 2 hemoglobin levels, and those missing baseline covariates.

### Exposure

The exposure of interest was the ART core drug class (NNRTI, INSTI or PI) prescribed as part of an ART regimen. Participants switching to different core drugs within the same class were considered to be continually exposed to the same core drug class. Individuals with a gap in ART use of 6 or more months were censored at the start of the gap and did not re-enter the study.

Person-time on INSTIs or PIs was compared to the reference of NNRTI use. In addition, some PLWH in this cohort had prescriptions for multiple core classes simultaneously. Participants with regimens containing more than 1 core class were categorized separately as users of “multiple core classes” in analyses. Boosting agents (e.g. boosted ritonavir, or cobicistat) were not considered a 2^nd^ core agent.

### Outcome ascertainment

Hb levels, expressed in grams per deciliter (g/dl), were ascertained using inpatient and outpatient laboratory data obtained as part of clinical care. Outcomes included incident anemia (first Hb measure below 10 g/dl), incident severe anemia (first Hb measure below 7.5 g/d) and changes in hemoglobin level. Another outcome, chronic anemia, defined as anemia lasting for ≥6 months was also examined. Chronic anemia was defined as Hb lab results on two separate occasions at least 6 months apart which were consistently in the anemic range without any Hb values above the anemia range during this 6-month period.

### Participant characteristics

Characteristics that were analyzed as confounders of the association between ART core drugs and incident anemia, severe anemia or change in hemoglobin overtime included: age, sex, race/ethnicity, CNICS site, hepatitis C virus (HCV) coinfection, kidney function measured using estimated glomerular filtration rate (eGFR, categorized as <30, 30-59, or ≥60 mL/minute/1.73 m^2^) [22], CD4 count (categorized as ≥500, 350-499, 200-399, 100-199 or <100 cells/mm^3^), viral load (VL, assessed as log_10_(VL+1)), and time in care at CNICS sites, defined as time from cohort entry date until the last available CNICS activity: either last lab date or last visit. HCV, eGFR, CD4 count and VL were assessed as part of clinical care visits and were time-updated as repeated measures occurred. All covariates were selected *a priori*, based on review of the literature and clinical knowledge. In addition, assessment of self-reported ART adherence was available for a subset of ~55% of the study population who were in care after each individual site initiated a clinical assessment of patient reported outcomes including adherence [23].

### Statistical analysis

Baseline characteristics are presented for all participants at the cohort entry date. Median and interquartile range (IQR) are displayed for continuous variables and frequencies and proportions are displayed for categorical variables.

Two multivariable Cox proportional hazards regression analyses were conducted, one among the subset of PLWH who were anemia-free at baseline to determine associations between time-updated NNRTI, PI, and INSTI use and development of anemia, and another among the subset of participants who were free of severe anemia at baseline to determine associations between time-updated NNRTI, PI, and INSTI use and development of severe anemia.

Participants were censored at a) the time they developed the outcome of interest, or b) at the time of last activity in CNICS, or c) at the time of death, or d) at the time of administrative censoring per site, or e) at the time they no longer were prescribed one of the ART core classes, whichever came first. The timescale for the models was time since cohort entry. Complete case analysis methods were used (<2% had missing data).

In a sensitivity analysis, we examined ART-naïve PLWH who initiated a drug in one of the ART classes of interest during study follow-up. Follow-up in this analysis began when a person began their initial ART regimen and extended until the earliest time of anemia occurrence, last activity in CNICS, time of death, administrative censoring, or at the time their initial regimen ended. PI or INSTI use were compared to the reference, NNRTI use. We also examined the change in Hb overtime using mixed models among this ART-naïve population.

Linear mixed effects models with random slopes for time were used to examine the association of ART core classes with Hb levels among all PLWH after adjustment for the same characteristics as in the incident anemia and severe anemia analyses. Mixed-effects models utilize random slopes and intercepts at the participant level to handle irregular patterns of repeated measures over follow-up [24]. All analyses were performed using Stata version 14.2.

### Patient and public involvement

There was no patient or public participation in the present study.

## Results

In total, 16,505 PLWH met inclusion criteria and were included in these analyses (Figure 1). Participants had an average of 11 outpatient hemoglobin values measured during a median follow-up of 4.9 (IQR 3.0-7.2) years. A total of 12,626 (76%) were free of anemia at baseline, and 15,357 (93%) were free of severe anemia at baseline. Table 1 provides baseline characteristics for all study participants, and additionally shows characteristics for the 1,040 participants who developed anemia during follow-up as well as the 488 participants who developed severe anemia during follow-up. The mean age of study participants was 46 years at cohort entry, 20% were female, and 19% were co-infected with HCV. At baseline, 18% were prescribed NNRTIs, 53% were prescribed PIs, 14% were prescribed INSTIs and 16% used multiple cores. INSTI core agents were increasingly used over the last few years of the enrollment period (Figure 2), and the proportion of those prescribed multiple cores were comprised of INSTI plus another core class with increasing frequency as study years progressed.

**Table 1.**
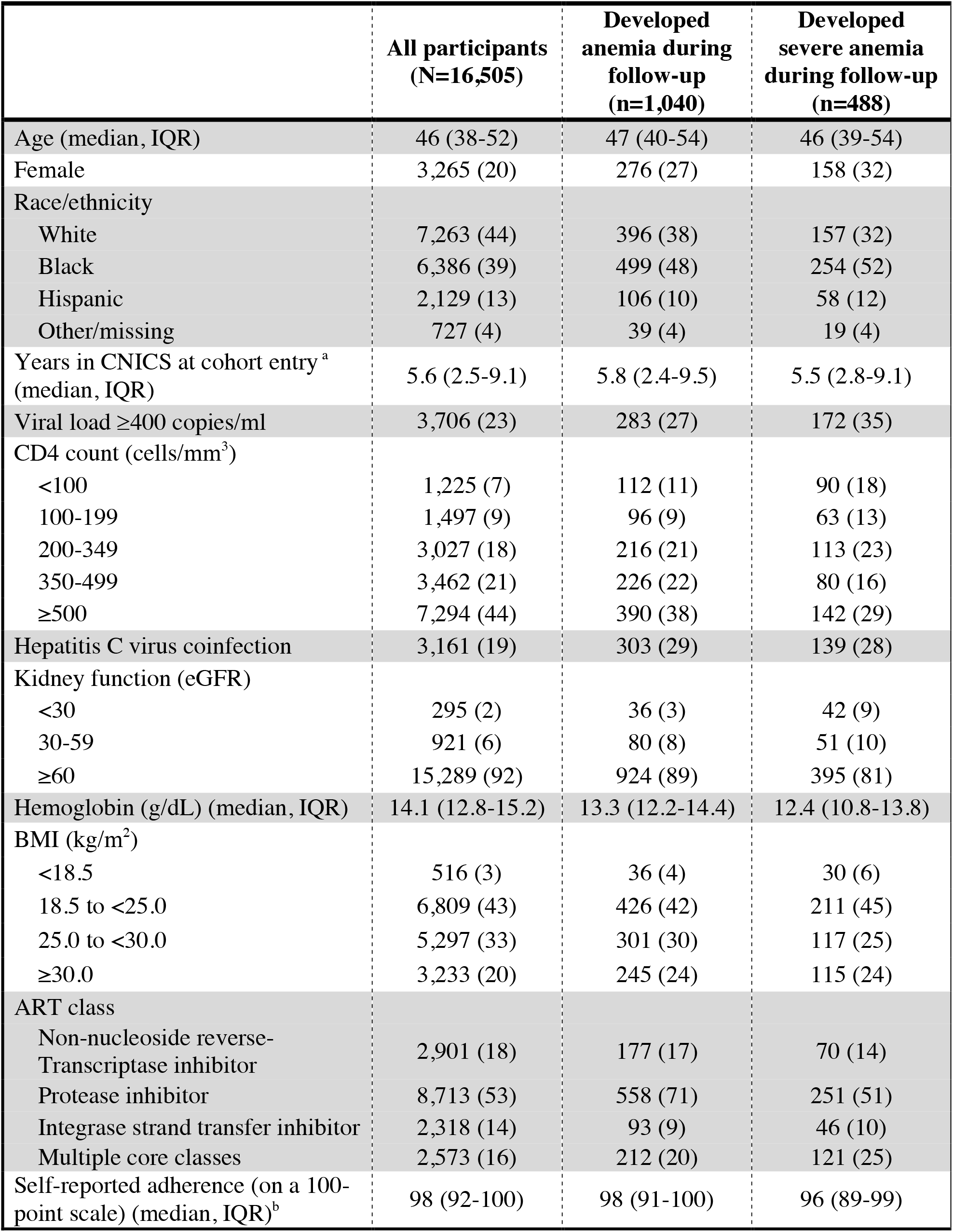
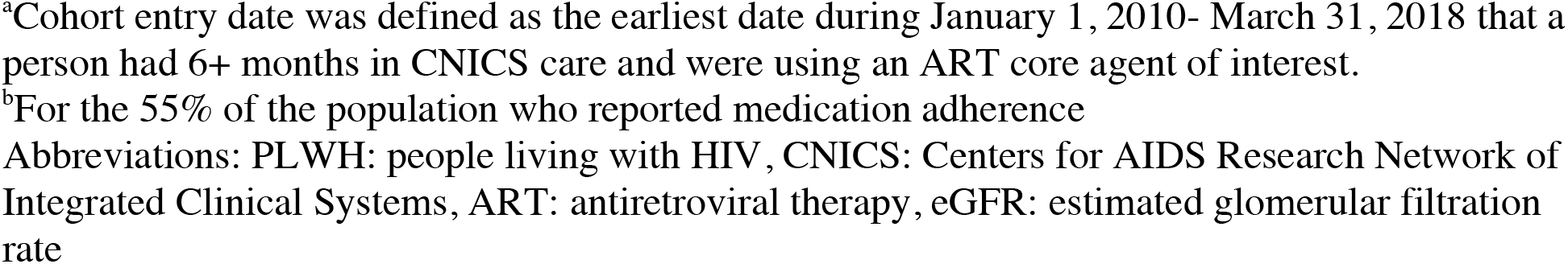
Baseline characteristics of PLWH in CNICS who were receiving an ART core agent of interest^a^

|  | **All participants**<br>**(N=16,505)** | **Developed**<br>**anemia during**<br>**follow-up**<br>**(n=1,040)** | **Developed**<br>**severe anemia**<br>**during follow-up**<br>**(n=488)** |
| --- | --- | --- | --- |
| Age (median, IQR) | 46 (38-52) | 47 (40-54) | 46 (39-54) |
| Female | 3,265 (20) | 276 (27) | 158 (32) |
| Race/ethnicity |  |  |  |
| White | 7,263 (44) | 396 (38) | 157 (32) |
| Black | 6,386 (39) | 499 (48) | 254 (52) |
| Hispanic | 2,129 (13) | 106 (10) | 58 (12) |
| Other/missing | 727 (4) | 39 (4) | 19 (4) |
| Years in CNICS at cohort entry<sup>a</sup><br>(median, IQR) | 5.6 (2.5-9.1) | 5.8 (2.4-9.5) | 5.5 (2.8-9.1) |
| Viral load $\geq 400$ copies/ml | 3,706 (23) | 283 (27) | 172 (35) |
| CD4 count (cells/mm<sup>3</sup>) |  |  |  |
| <100 | 1,225 (7) | 112 (11) | 90 (18) |
| 100-199 | 1,497 (9) | 96 (9) | 63 (13) |
| 200-349 | 3,027 (18) | 216 (21) | 113 (23) |
| 350-499 | 3,462 (21) | 226 (22) | 80 (16) |
| $\geq 500$ | 7,294 (44) | 390 (38) | 142 (29) |
| Hepatitis C virus coinfection | 3,161 (19) | 303 (29) | 139 (28) |
| Kidney function (eGFR) |  |  |  |
| <30 | 295 (2) | 36 (3) | 42 (9) |
| 30-59 | 921 (6) | 80 (8) | 51 (10) |
| $\geq 60$ | 15,289 (92) | 924 (89) | 395 (81) |
| Hemoglobin (g/dL) (median, IQR) | 14.1 (12.8-15.2) | 13.3 (12.2-14.4) | 12.4 (10.8-13.8) |
| BMI (kg/m<sup>2</sup>) |  |  |  |
| <18.5 | 516 (3) | 36 (4) | 30 (6) |
| 18.5 to <25.0 | 6,809 (43) | 426 (42) | 211 (45) |
| 25.0 to <30.0 | 5,297 (33) | 301 (30) | 117 (25) |
| $\geq 30.0$ | 3,233 (20) | 245 (24) | 115 (24) |
| ART class |  |  |  |
| Non-nucleoside reverse-Transcriptase inhibitor | 2,901 (18) | 177 (17) | 70 (14) |
| Protease inhibitor | 8,713 (53) | 558 (71) | 251 (51) |
| Integrase strand transfer inhibitor | 2,318 (14) | 93 (9) | 46 (10) |
| Multiple core classes | 2,573 (16) | 212 (20) | 121 (25) |
| Self-reported adherence (on a 100-point scale) (median, IQR)<sup>b</sup> | 98 (92-100) | 98 (91-100) | 96 (89-99) |

<sup>a</sup>Cohort entry date was defined as the earliest date during January 1, 2010- March 31, 2018 that a person had 6+ months in CNICS care and were using an ART core agent of interest.
<sup>b</sup>For the 55% of the population who reported medication adherence
Abbreviations: PLWH: people living with HIV, CNICS: Centers for AIDS Research Network of Integrated Clinical Systems, ART: antiretroviral therapy, eGFR: estimated glomerular filtration rate

**Figure 2.**
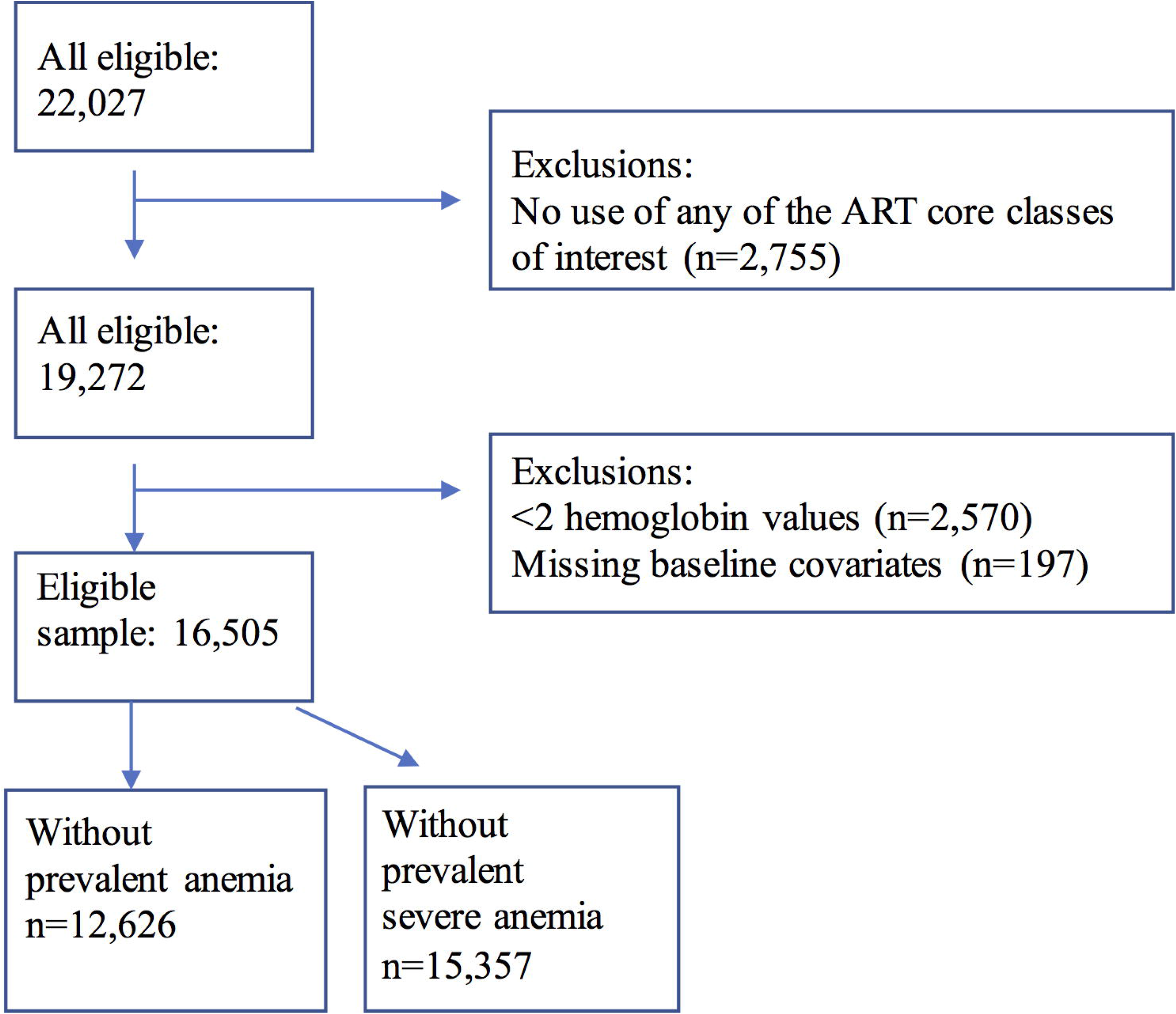
Proportion of study population using various ART classes during complete years of study follow-up. This figure shows the trends in use of the ART core classes during 2010-2017.

The incidence of anemia was 2.1/100 person-years and the incidence of severe anemia was 0.8/100 person-years. INSTI use was associated with an increased risk of anemia (aHR 1.17, 95%CI 0.94-1.47) compared to NNRTIs, though this was not statistically significant (Table 2). Use of multiple core classes together was also associated with an increased risk of anemia (aHR 1.30, 95%CI 1.06-1.60)) while no associations were found between PI use and anemia (aHR of 1.00, 95%CI 0.83-1.21). In adjusted analyses restricted to participants free of severe anemia at baseline (Table 2), INSTI use was associated with an increased risk of severe anemia (aHR 1.55, 95%CI 1.11-2.17) compared to NNRTI use. Time on multiple ART core classes was associated with an increased risk of anemia (aHR 1.35 (0.99-1.85), though this was not statistically significant, and no association was found for PIs (aHR 1.01 (0.74-1.36) Among the 12,626 PLWH who were free of anemia at baseline, 225 developed chronic anemia (lasting for ≥6 months), during follow-up. For chronic anemia, results were similar to those in the primary analysis. Relative to NNRTI use, person-time on multiple core classes was associated with an aHR 2.26 (95%CI 0.98-5.23), person-time on an INSTI with an aHR of 1.94 (95%CI 0.80-4.68) and person-time on a PI with an aHR of 1.29 (95%CI 0.55-3.06).

**Table 2:** Association of ART classes with incident anemia (hemoglobin<10 g/dL), severe anemia (hemoglobin<7.5 g/dL) or chronic anemia (6+ months of anemia)

| ART Regimen | Hazard Ratio of incident anemia (n=12,626) |  | Hazard Ratio of incident severe anemia (n=15,357) |  | Hazard Ratio of incident chronic anemia (n=12,626) |  |
| --- | --- | --- | --- | --- | --- | --- |
|  | Unadjusted | Adjusted<sup>a</sup> | Unadjusted | Adjusted<sup>a</sup> | Unadjusted | Adjusted<sup>a</sup> |
| NNRTI (REF) | 1.00 | 1.00 | 1.00 | 1.00 | 1.00 | 1.00 |
| PI | 1.26 (1.05-1.52) | 1.00 (0.83-1.21) | 1.37 (1.02-1.83) | 1.01 (0.74-1.36) | 1.43 (0.61-3.35) | 1.29 (0.55-3.06) |
| INSTI | 1.39 (1.11-1.75) | 1.17 (0.94-1.47) | 1.96 (1.40-2.75) | 1.55 (1.11-2.17) | 2.05 (0.85-4.94) | 1.94 (0.80-4.68) |
| Multiple core classes | 2.02 (1.65-3.48) | 1.30 (1.06-1.60) | 2.70 (1.99-3.67) | 1.35 (0.99-1.85) | 3.46 (1.51-7.93) | 2.26 (0.98-5.23) |

<sup>a</sup>Adjusted for age, sex, race/ethnicity, CNICS site, Hepatitis C virus status, CD4 count, viral load, kidney function (eGF)
Abbreviations: ART: antiretroviral therapy, NNRTI: non-nucleoside reverse-transcriptase inhibitor, PI: protease inhibitor, INSTI: integrase strand transfer inhibitor

Average hemoglobin levels remained steady during follow-up; the mean level was 14.1 g/dL (IQR 12.7-15.1) at baseline and 14.0 (IQR 12.6-15.2) g/dL at the last available measurement per person. Relative to NNRTI use, a decrease in hemoglobin level over time was associated with both INSTI use (−0.06 g/dL per year, 95%CI −0.10, −0.03) and use of multiple core classes (−0.14, 95%CI −0.18, −0.11). No association was found for PI use (−0.01, 95%CI − 0.04, 0.03). (Table 3).

**Table 3.** Association of ART classes with change in hemoglobin level during follow-up in adjusted analyses (linear mixed-effect model); N=16,505

| ART class | Coefficient<sup>a</sup> | 95% CI | P-value |
| --- | --- | --- | --- |
| NNRTI (REF) |  |  |  |
| PI | -0.01 | -0.04, 0.03 | 0.675 |
| INSTI | -0.06 | -0.10, -0.03 | <0.001 |
| Multiple core classes | -0.14 | -0.18, -0.11 | <0.001 |

<sup>a</sup>Coefficient is the mean difference per year in Hb (g/dL) for each core regimen relative to the NNRTI core regimen, after adjustment for site, age, sex, race/ethnicity, hepatitis C virus coinfection, CD4 cell count, viral load, eGFR and years in study.
Abbreviations: ART: antiretroviral therapy, NNRTI: non-nucleoside reverse-transcriptase inhibitor, PI: protease inhibitor, INSTI: integrase strand transfer inhibitor

The sensitivity analysis restricted to ART-naïve PLWH included 6,426 PLWH who were free of prevalent anemia at baseline, of whom 378 developed anemia. Compared to NNRTI initiators, those initiating a PI had an aHR of 0.78 (0.56-1.08) while those initiating an INSTI had an aHR of 1.15 (95%CI 0.92-1.45) (Supplemental Table 1). The mixed model examining change in Hb overtime among ART-naïve PLWH initiating one of the ART core classes of interest included 7,264 participants. Compared to NNRTI initiators, a decrease in Hb was found for PI use (−0.8g/dL per year, 95%CI −0.16, −0.01), while INSTI use was associated with a larger decrease in Hb level overtime (−0.15g/dL per year, 95%CI −0.22, −0.09) (Supplemental Table 2).

## Discussion

In this study of 16,505 PLWH in care within the United States in the current treatment era (2010 and after), we observed that INSTI use, or time on multiple core ART classes used was associated with decreases in hemoglobin levels during follow-up compared to NNRTI use. We found that INSTI use was associated with an elevated aHR for anemia (aHR 1.17, 95%CI 0.94-1.47) and a significantly elevated aHR for severe anemia (1.55, 95%CI 1.11-2.17) as well as a significant decrease in hemoglobin levels over time. Furthermore, the naïve user analysis had nearly identical findings, although not significant with larger CIs. These findings could have implications for the treatment approach that should be used in people with risk factors for anemia.

This study’s strengths include its large and geographically diverse study population and longitudinal data structure. However, there are limitations of this study to consider including the observational nature of the data, which may be subject to residual confounding. Additionally, we did not exclude participants with conditions strongly associated with anemia or hemoglobin level through non-traditional HIV mechanisms, including those on dialysis, receiving erythropoietin, or with severe bleeding, which likely caused some of the anemia cases in this analysis. However, in the sensitivity analysis focusing on factors associated with chronic anemia (less likely due to bleeding), findings for INSTI vs. NNRTI core regimens were similar to those including all PLWH who became anemic. A final limitation is that ART medication use comes from prescriptions written which may not be filled or may be filled but never taken, although self-reported adherence was high (approximately 98%) in the subset for whom adherence information was available. CNICS participants who provided adherence information have been shown to be representative of the overall population of PLWH in CNICS [23, 25]. Finally, the fact that this study was conducted among PLWH in care in the U.S. who are on ART may limit the generalizability of findings to PLWH who are not on ART or who live outside of the U.S.

One study, conducted during the newer-era of HIV treatment with drugs other than AZT, (during 2008-2012) presented findings for AZT versus non-AZT regimens, finding an increased risk of anemia among AZT compared to non-AZT regimens (HR=2.84, 95%CI 1.52-5.31) [26]. However, anemia risk was not analyzed separately for the use of specific classes of ART, resulting in the inability of comparison to the present study’s findings and a lack of generalizability to PLWH who are treated with newer ART core agents.

It is possible that PLWH in our study whose HIV is progressing due to resistance or other complications may get switched to an INSTI. This, in addition to prior knowledge that poorly controlled HIV parameters are on their own a risk factor for anemia [6, 27, 28], could result in confounding by indication. However, the switch to INSTI core regimens since their approval in 2007 has been widespread in this population (Figure 2) and INSTIs are recommended for use as initial regimens [29]. In addition, we rigorously controlled for many of the important HIV-related factors that correspond to poorly-controlled HIV, and our sensitivity analysis examining new users of ART medications failed to reinforce the notion that an increased risk of anemia among INSTI core regimen users could be entirely explained by sicker participants getting switched to these therapies.

PLWH on multiple core classes were in a different category in our analyses. There are several reasons PLWH may be prescribed multiple core classes. For example, sometimes PLWH are prescribed multiple core classes to ensure they receive a complete regimen while awaiting approval for specific agents from their insurance company. However, the primary concern was that they were receiving multiple core classes due to provider concerns such as prior failed regimens which may also increase their risk of anemia.

In conclusion, in this large, diverse, multicenter cohort of PLWH, we found that INSTI use and time on multiple ART core classes were associated with progression to anemia and a lowering of Hb level. INSTI use was also associated with severe anemia risk. Our findings suggest that careful selection of ART regimen could mitigate anemia development, although this anemia risk needs to be balanced with the possibility of improvement in overall HIV care [30]. Further research is needed to replicate the finding of INSTI core regimen use and anemia risk and to understand the underlying mechanisms. If confirmed, screening for anemia development in users of INSTIs may be beneficial.

