## Supplemental Table 1 for "Class of antiretroviral drugs and anemia risk in the current treatment era"

Supplemental Table 1: Association of ART classes with incident anemia among naïve users

| **ART class** | | **Hazard Ratio of incident anemia (n=6,426)** | |
| --- | --- | --- | --- |
|  | **Unadjusted** | | **Adjusted^a^** |
| NNRTI (REF) | 1.00 | | 1.00 |
| PI | 0.92 (0.67-1.27) | | 0.78 (0.56-1.08) |
| INSTI | 1.73 (1.39-2.15) | | 1.15 (0.92-1.45) |

Abbreviations: ART: antiretroviral therapy, NNRTI: non-nucleoside reverse-transcriptase inhibitor, PI: protease inhibitor, INSTI: integrase strand transfer inhibitor
