## Supplemental Table 2 for "Class of antiretroviral drugs and anemia risk in the current treatment era"

Supplemental Table 2. Association of ART classes with change in hemoglobin level during follow-up in adjusted analyses among naïve users (linear mixed-effect model); N=7,264

| **ART class** | **Coefficient^a^** | **95% CI** | **P-value** |
| --- | --- | --- | --- |
| NNRTI (REF) |  |  |  |
| PI | -0.08 | -0.16, -0.01 | 0.031 |
| INSTI | -0.15 | -0.22, -0.09 | <0.001 |

^a^Coefficient is the mean difference per year in Hb (g/dL) for each core regimen relative to the NNRTI core regimen, after adjustment for site, age, sex, race/ethnicity, hepatitis C virus coinfection, CD4 cell count, viral load, eGFR and years in study.

Abbreviations: ART: antiretroviral therapy, NNRTI: non-nucleoside reverse-transcriptase inhibitor, PI: protease inhibitor, INSTI: integrase strand transfer inhibitor
